# *N*^6^-methyladenosine binding proteins negatively regulate HIV-1 infectivity and viral production

**DOI:** 10.1101/257410

**Authors:** Wuxun Lu, Nagaraja Tirumuru, Pratibha C. Koneru, Chang Liu, Mamuka Kvaratskhelia, Chuan He, Li Wu

**Affiliations:** Center for Retrovirus Research, Department of Veterinary Biosciences, The Ohio State University, Columbus, Ohio 43210, USA; Division of Infectious Diseases, University of Colorado School of Medicine, Aurora, Colorado 80045, USA; Department of Chemistry, Department of Biochemistry and Molecular Biology, Institute for Biophysical Dynamics, The University of Chicago, Chicago, Illinois 60637, USA; Howard Hughes Medical Institute, The University of Chicago, Chicago, Illinois 60637, USA

**Keywords:** HIV-1 infection, HIV-1 production, RNA m^6^A modification, m^6^A-binding proteins

## Abstract

**Background:** The internal *N*^6^-methyladenosine (m^6^A) modification of cellular mRNA regulates post-transcriptional gene expression. The YTH domain family proteins (YTHDF1-3, or Y1-3) bind to m^6^A-modified cellular mRNA and modulate its metabolism and processing, thereby affecting protein translation in cells. We previously reported that HIV-1 RNA contains m^6^A modification and that Y1-3 proteins inhibit HIV-1 infection by decreasing HIV-1 reverse transcription. Here we extended our studies to further understand the mechanisms of Y1-3-mediated inhibition of HIV-1 infection and viral production.

**Results:** Overexpression of Y1-3 proteins in HIV-1 target cells decreased viral genome RNA (gRNA) levels and inhibited early and late reverse transcription. Purified recombinant Y1-3 proteins preferentially bound to the m^6^A-modified 5’ leader sequence of gRNA compared with its unmodified RNA counterpart, consistent with the strong binding of Y1-3 to HIV-1 gRNA in infected cells. HIV-1 mutants with two altered m^6^A modification sites in the 5’ leader sequence of gRNA demonstrated significantly lower infectivity compared with wild-type HIV-1, confirming that these sites are important for viral infection. HIV-1 produced from cells with knockdown of endogenous Y1, Y3, or Y1-3 proteins together showed increased viral infectivity compared with HIV-1 produced from control cells. Interestingly, we found that Y1-3 proteins and HIV-1 Gag formed a complex with RNA in HIV-1-infected target cells.

**Conclusions:** Our results suggest the inhibitory effects of Y1-3 proteins on HIV-1 infection and provide new insight into the mechanisms of m^6^A modification of HIV-1 RNA in regulating viral replication, which clarify some discrepancies in the previously published studies in this area.

## Background

Among the more than 100 distinct modifications identified in mRNAs in different organisms, *N*^6^-methyladenosine (m^6^A) methylation is the most prevalent internal modification, accounting for 0.1% of adenosines in mammalian mRNAs [1]. The dynamic addition, removal, and recognition of m^6^A in cellular RNAs are coordinately regulated by three groups of host proteins, including methyltransferases (termed writers), demethylases (erasers), and m^6^A-binding proteins (readers). The writers include methyltransferase-like 3 (METTL3), methyltransferase-like 14 (METTL14), and Wilms tumor 1-associated protein (WTAP), while erasers include fat mass and obesity-associated protein (FTO) and alkB homologue 5 (ALKBH5) [2]. The readers are YT521-B homology (YTH) domain family proteins (YTHDF1, YTHDF2, YTHDF3 and YTHDC1) that specifically recognize m^6^A modification via a conserved m^6^A-binding pocket in the YTH domain [3-5]. These reader proteins modulate m^6^A-modifed mRNA stability and translation, therefore playing an important role in modulating post-transcriptional gene expression [6-10].

Three recent studies highlighted the importance of m^6^A modifications of HIV-1 RNA in regulating viral replication and gene expression [11-13]. Despite some consistent results, there are discrepancies in the locations, effects, and mechanisms of m^6^A modification of HIV-1 RNA in these studies [11-13]. Our published study identified m^6^A modifications in the 5’ and 3’ untranslated regions (UTRs), as well as in the *rev* and *gag* genes of the HIV-1 genome [13]. We previously reported that overexpression of Y1-3 proteins in cells inhibits HIV-1 infection by primarily decreasing HIV-1 reverse transcription, while knockdown of endogenous Y1-3 in Jurkat CD4^+^ T-cell line or primary CD4^+^ T-cells increases HIV-1 infection [13]. However, the underlying mechanisms of Y1-3-mediated inhibition of HIV-1 infection remain unclear.

Here we report that Y1-3 inhibited HIV-1 infection in target cells by lowering viral genome RNA (gRNA) levels and reverse transcription products. We demonstrate that the m^6^A-modified HIV-1 RNA fragment preferentially bound to Y1-3 proteins compared with an un-modified RNA counterpart, and mutations of two m^6^A sites in the 5’ UTR significantly decreased viral infectivity. Knockdown of endogenous Y1, Y3 or Y1-3 proteins together in virus-producing cells positively modulated HIV-1 infectivity. Furthermore, we found that endogenous Y1-3 proteins and HIV-1 Gag protein formed a complex with RNAs in HIV-1-infected target cells. Together, these findings suggest new mechanisms by which Y1-3 proteins mediate HIV-1 inhibition during early steps of the viral life cycle.

## Results

### Overexpression of Y1-3 proteins in target cells inhibits post-entry HIV-1 infection by decreasing HIV-1 gRNA level and inhibiting viral reverse transcription

Our previous study showed that Y1-3 proteins negatively regulate HIV-1 post-entry infection in target cells, including primary CD4^+^ T-cells [13]. To better understand the underlying mechanisms, we compared HIV-1 gRNA levels, early and late reverse transcription (RT) products in cells overexpressing individual Y1-3 proteins and control cells after HIV-1 infection. We first used vesicular stomatitis virus glycoprotein (VSV-G)-pseudotyped single-cycle HIV-1 expressing firefly luciferase to infect HeLa cells stably expressing each individual Y1-3 proteins (HeLa/Y1-3) or control cells transduced with an empty vector (HeLa/Vector). Consistent with our previous results [13], Y1-3 overexpression (Fig. 1A) resulted in significantly lower levels of HIV-1 post-entry infection (26%, 8.6%, and 23% in HeLa/Y1, HeLa/Y2 and HeLa/Y3 cells, respectively) compared with HeLa/Vector control cells (set as 100%, Fig. 1B).

**Fig. 1.**
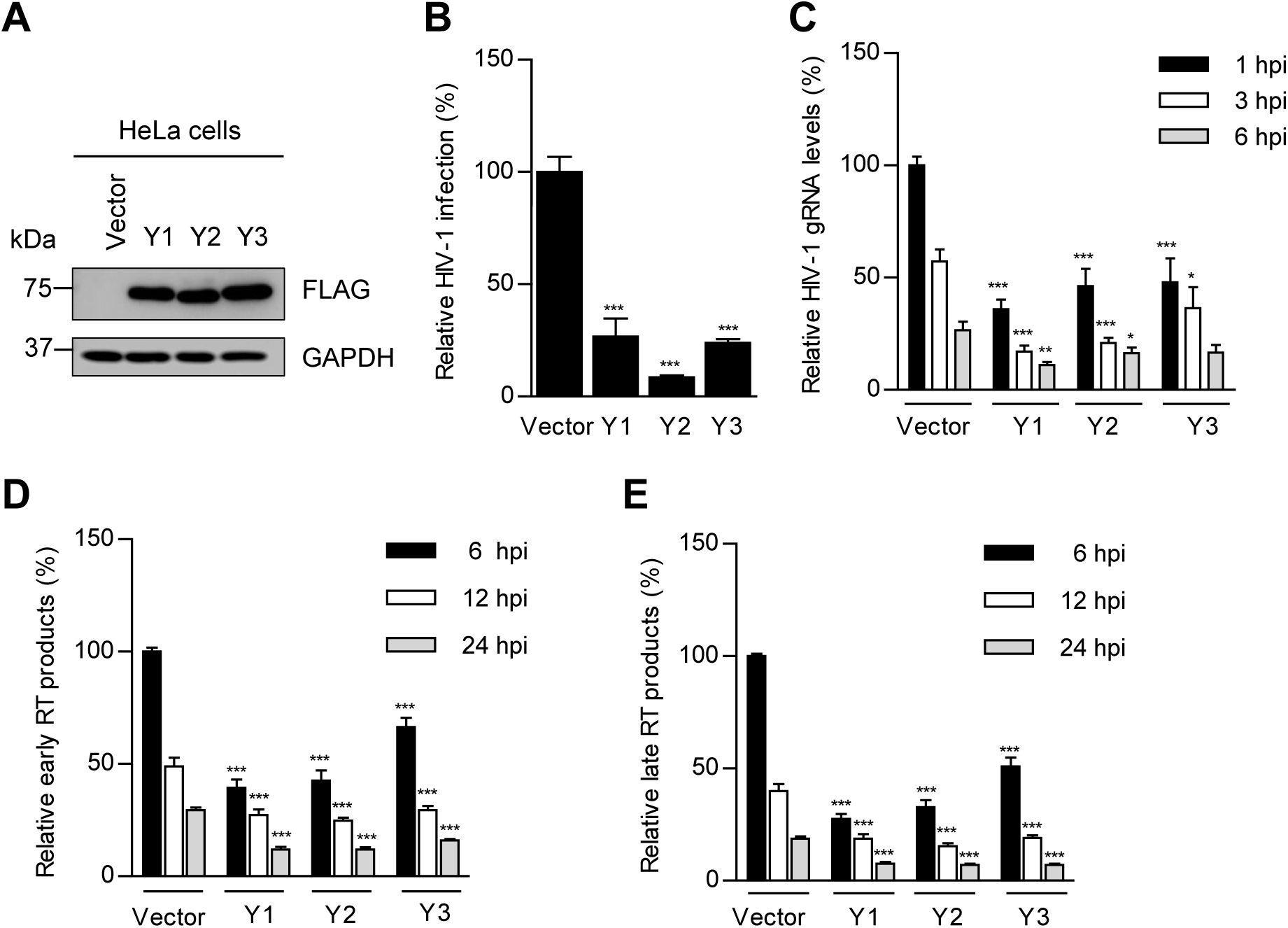
Overexpression of Y1-3 proteins inhibits HIV-1 infection through decreasing HIV-1 gRNA and viral reverse transcription. **(A)** Overexpression of Y1-3 proteins in HeLa/Y1-3 cell lines. **(B)** Cells were infected with HIV-1 Luc/VSV-G (MOI=1), and viral infection was measured by luciferase activity at 24 h post-infection (hpi) (n=3 independent experiments, same in the following). **(C)** Quantified HIV-1 gRNA levels after infection of HeLa/Y1-3 cells and vector control cells (n=3). **(D)** Early reverse transcription (RT) product levels at 6, 12 and 24 hpi (n=3). **(E)** Late RT product levels at 6, 12 and 24 hpi (n=3). Results are shown as mean ± SEM. Dunnett’s multiple comparison test was used to determine statistical significance. * P < 0.05, *** P < 0.0005, compared each group with vector control cells in each corresponding experiment. Data are from at least 3 independent experiments with biological duplicates.

To examine whether Y1-3 proteins alter the levels of HIV-1 gRNA in the infected cells, HeLa/Y1-3 or control cells were infected with single-cycle HIV-1 and total RNAs from the cells were isolated at 1, 3, and 6 h post-infection (hpi). The levels of HIV-1 gRNA were quantified using RT-qPCR [14]. At each time point tested, HIV-1 gRNA levels were lower in HeLa/Y1-3 relative to those in HeLa/Vector control cells, and HIV-1 gRNA level gradually declined after infection as expected (Fig. 1C). These results indicate that overexpression of Y1-3 proteins leads reduced levels of HIV-1 gRNA, likely by decreasing viral RNA stability.

Our previous study showed that Y1-3 proteins inhibit accumulation of HIV-1 late RT products in infected cells [13]; however, it is unclear whether Y1-3 proteins affect HIV-1 early RT efficiency. To address this question, HeLa/Y1-3 cells or HeLa/Vector control cells infected by HIV-1-Luc/VSV-G were collected at 6, 12 and 24 hpi for quantification of HIV-1 early and late RT products by quantitative PCR (qPCR) [15]. At each time point, the levels of both early and late RT products were significantly lower in HeLa/Y1-3 cells compared with vector control cells (Fig. 1D and 1E). Compared with vector control cells at 6 hpi, Y1-3 overexpression decreased HIV-1 gRNA levels, and further decreased early and late RT products (Fig. 1C-E), suggesting that overexpression of Y1-3 proteins decreases viral gRNA stability and inhibits early RT product synthesis. At 24 hpi, the levels of early and late RT products decreased to 37-54%, while HIV-1 infection decreased to 8.6-26% in HeLa/Y1-3 cells compared with vector control cells (set to 100% in Fig. 1B, 1D and 1E). Together, these results suggest that overexpression of Y1-3 proteins in target cells inhibits HIV-1 infection mainly by decreasing HIV-1 gRNA level and inhibiting viral reverse transcription.

### Overexpression of Y1-3 proteins in HeLa/CD4 target cells inhibits wild-type (WT) HIV-1 replication

Since the *firefly luciferase* gene in HIV-1-Luc/VSV-G virus has consensus sequences recognized by m^6^A writers, its mRNA may also have m^6^A modifications. To exclude the effects of m^6^A modification of *luciferase* mRNA on HIV-1 infection, we used WT, replication-competent HIV-1_NL4-3_ to infect HeLa cells overexpressing CD4 and individual Y1-3 proteins (HeLa/CD4/Y1-3) or vector control cells. Using flow cytometry, we confirmed that the majority of these cells were double positive for the HIV-1 primary receptor CD4 and co-receptor CXCR4 (71-81%), which would allow efficient fusion-mediated viral entry (Fig. 2A). Immunoblotting results also confirm that HeLa/CD4/Y1-3 cells stably expressed FLAG-tagged individual Y1-3 proteins (Fig. 2B).

**Fig. 2.**
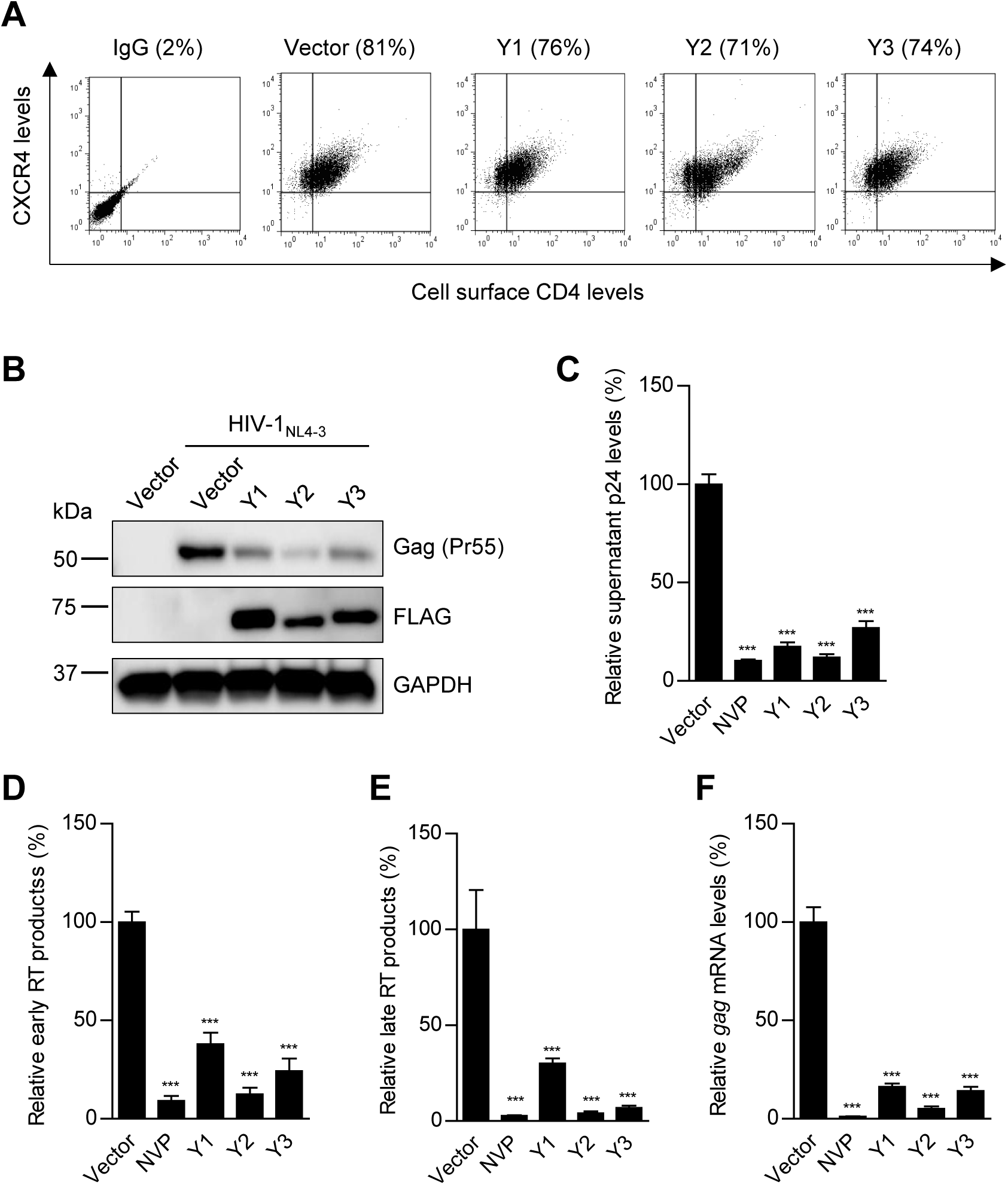
Overexpression of Y1-3 proteins in HeLa/CD4 target cells inhibits wild-type HIV-1 replication. **(A)** Surface levels of CD4 (exogenous) and CXCR4 (endogenous) in HeLa/CD4 cells overexpressing individual Y1-3 proteins or vector control cells were analyzed by flow cytometry. Isotype-matched IgG was used as a negative control for immunostaining. The numbers on the top of the plots indicate the percentages of CD4-and CXCR4-positive cells. **(B-F)** HeLa/CD4 cells overexpressing individual Y1-3 proteins were infected with HIV-1_NL4-3_ (MOI=1) for 72 h. **(B)** HIV-1 Gag protein expression and over-expression of FLAG-tagged Y1-3 proteins in HeLa/CD4 cells were confirmed by immunoblotting. **(C)** ELISA quantification of HIV-1 p24 levels in the supernatants from infected cells. **(D** and **E)** qPCR quantification of the levels of HIV-1 early RT products (D) and late RT products (E). **(F)** RT-qPCR quantification of HIV-1 *gag* mRNA in the cells. (C-F) The reverse transcriptase inhibitor nevirapine (NVP) was used to treat vector cells as a negative control for HIV-1 infection. All results are shown as mean ± SEM (n=3) and data presented are representative of 3 independent experiments. Dunnett’s multiple comparison test was used to determine statistical significance. ** P <0.005 and *** P <0.0005 compared with vector control cells.

In order to examine the effects of Y1-3 overexpression on HIV-1 replication in the full viral lifecycle, we collected infected cells or supernatants at 72 hpi for further analyses (Fig. 2B-F). Consistent with the results from single-cycle HIV-1-Luc/VSV-G (Fig. 1B), Y1-3 overexpression efficiently inhibited infection of WT HIV-1_NL4-3_ as Gag protein levels in cells and p24 (capsid) levels in supernatants were significantly lower in HeLa/CD4/Y1-3 cells compared with vector control cells (Fig. 2B and 2C). These results confirmed inhibitory effects of Y1-3 proteins on HIV-1 infection, and suggested that the inserted *luciferase* gene in HIV-1-Luc/VSV-G does not affect the observed phenotype. Infection of HeLa cells overexpressing Y1-3 proteins with WT HIV-1 also resulted in significantly lower early and late HIV-1 RT products (Fig. 2D and 2E, respectively), and consequently reduced *gag* mRNA levels compared with vector control cells (Fig. 2F). These data suggest that Y1-3 proteins inhibit WT HIV-1_NL4-3_ infection before or during the RT stage.

### Y1-3 proteins specifically bind to HIV-1 gRNA in infected HeLa/CD4 cells

To confirm Y1-3 binding to HIV-1 gRNA in infected cells, we performed immunoprecipitation (IP) of Y1-3 in HeLa/CD4/Y1-3 cell lines and vector control cells infected with WT HIV-1_NL4-3_ (Fig. 3A). To validate the specificity of Y1-3 binding to HIV-1 RNA, we also included a Y1-3 unrelated cellular protein MAL (<u>M</u>yD88 <u>a</u>dapter-like, also known as Toll/IL-1 receptor (TIR) domain-containing adapter protein, or TIRAP) [16, 17] as an additional negative control in the IP assay (Fig. 3A). We then quantified the amounts of Y1-3-bound HIV-1 gRNA using RT-qPCR assays [13]. We observed that Y1-3 specifically and efficiently bound to WT HIV-1 gRNA in HIV-1_NL4-3_ infected HeLa/CD4/Y1-3 cells compared with vector control or MAL-expressing cells (Fig. 3B). The levels of HIV-1 gRNA bound to Y1 appeared higher than those bound to Y2 and Y3 (Fig. 3B), which could be due to the higher expression of Y1 in HeLa/CD4 cells relative to Y2 and Y3 proteins (Fig. 3A) and consequently a higher level of IP products (Fig. 3B). These data confirm that Y1-3 proteins specifically bind to WT HIV-1_NL4-3_ gRNA in infected cells.

**Fig. 3.**
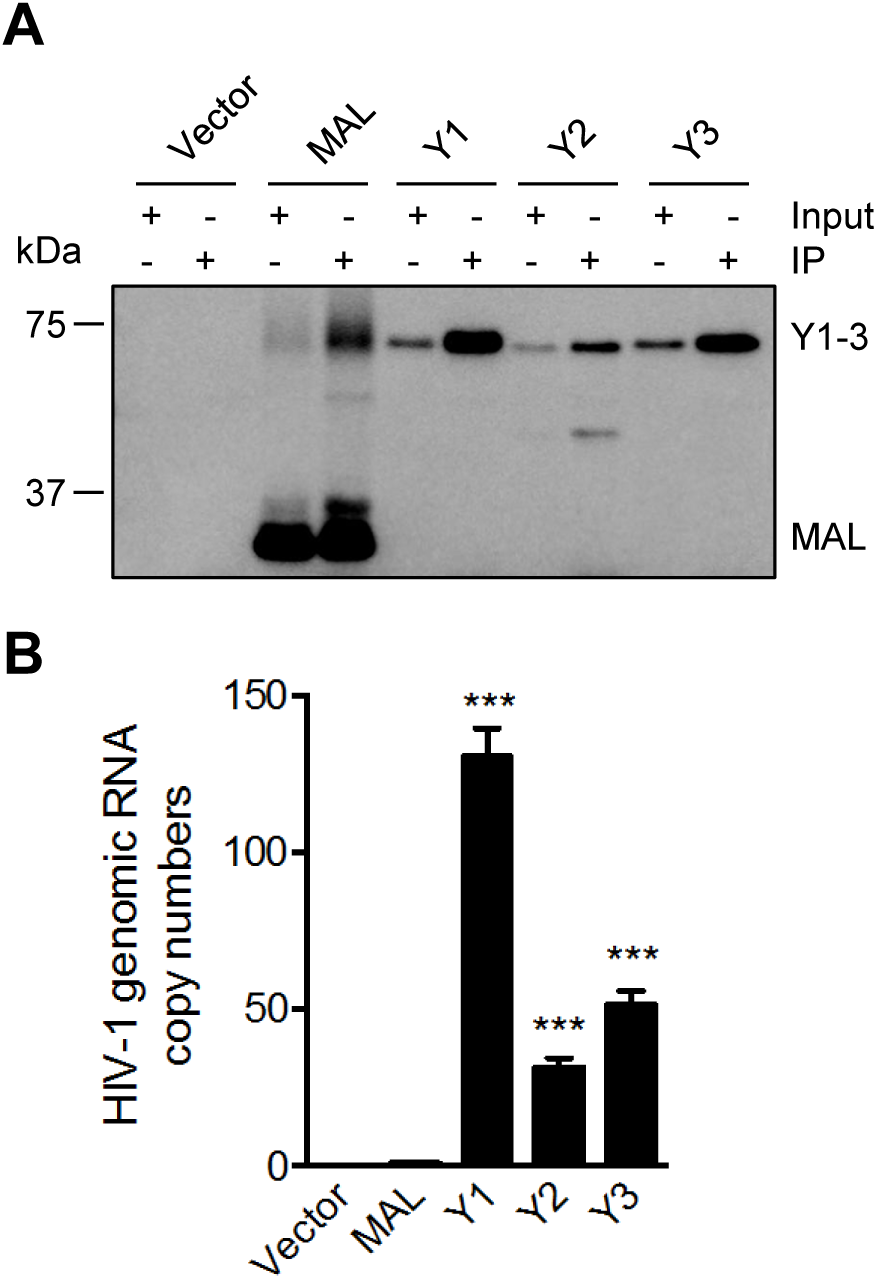
Y1-3 proteins specifically bind to HIV-1 gRNA in infected HeLa/CD4 cells. (**A**) Immunoblotting of Y1-3 in the input and immunoprecipitation (IP) samples from HeLa/CD4 cells infected with HIV-1. HeLa/CD4 cells stably overexpressing FLAG-tagged Y1-3 proteins, MAL (MyD88 adapter-like protein), or empty vector control cells were infected with HIV-1_NL4-3_ at an MOI of 5 for 3 h. FLAG antibodies were used to immunoprecipitate Y1-3 proteins or MAL in HeLa/CD4 cells after HIV-1 infection. (**B**) HIV-1 gRNA is bound by Y1-3 proteins expressed in HeLa/CD4 cells. HIV-1 infection of HeLa/CD4 cells overexpressing Y1-3 proteins, MAL, or empty vector control cells as described above in (A). Cell lysates were immunoprecipitated with anti-FLAG antibody, and HIV-1 *gag* RNA levels were quantified by RT-qPCR. Dunnett’s multiple comparison test was used to determine statistical significance. *** P <0.0005 compared with the vector control cells. Data presented are representative of 4 independent experiments.

### Purified recombinant Y1-3 proteins preferentially bind to an m^6^A-modifed HIV-1 RNA fragment *in vitro*

Our previous study showed that HIV-1 RNA contains m^6^A modifications at both the 5’ and 3’ UTR [13]. Given the critical role of the 5’ UTR in initiation of HIV-1 reverse transcription, in this study we focused on the m^6^A sites in the 5’ UTR of HIV-1 gRNA. The GGACU motif is the most predominant sequence for m^6^A modification [18, 19]. The m^6^A peak detected by high-throughput RNA sequencing in the 5’ UTR of HIV-1 gRNA harbors two GGACU motifs [13]. The first one is located in the primer-binding site (PBS), and the second is in upstream region of the dimer initiation sequence (DIS) (Fig. 4A). These two GGACU motifs overlap with m^6^A modifications in HIV-1 gRNA and are close to Y1-3 protein-binding peaks identified in the 5’ UTR of HIV-1 gRNA [13].

**Fig. 4.**
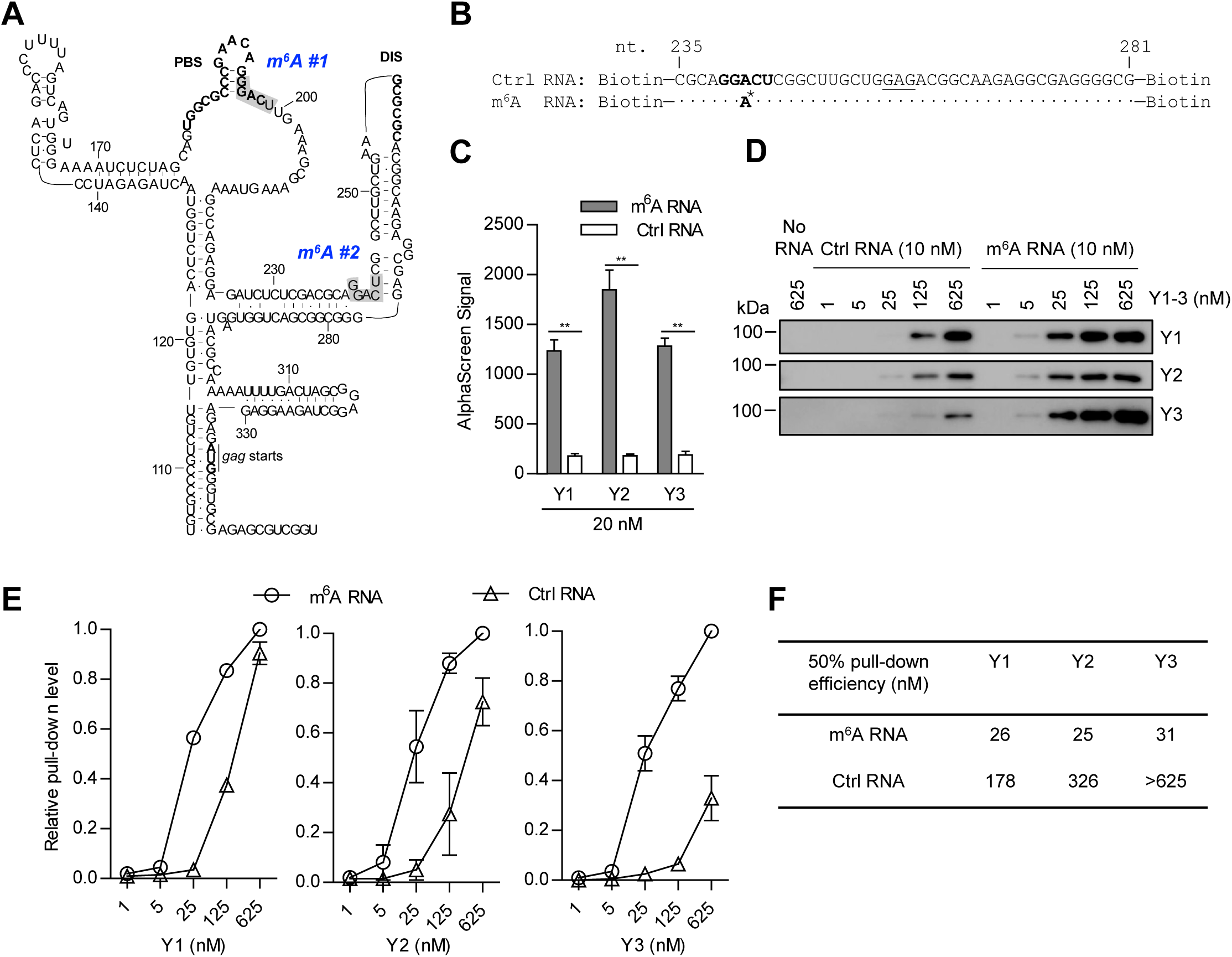
Purified recombinant Y1-3 proteins preferentially bind to an m^6^A-modifed HIV-1 RNA fragment *in vitro*. **(A)** Structure of partial HIV-1 RNA 5’ UTR containing two GGACU motifs (in gray background) that are labeled with blue words *m*^*6*^*A #1* (nt. 195-199) and *m*^*6*^*A #2* (nt. 239-243). PBS and DIS are highlighted in bold. The figure is adapted from Kharytonchyk et al. [24]. **(B)** Sequences of synthesized control (Ctrl) RNA and m^6^A-modified (m^6^A RNA) fragment containing the second GGACU motif (in bold) in HIV-1 RNA 5’ UTR corresponding to nt. 235-281. The DIS-containing sequence (AA**GCGCGC**) was replaced with nucleotides GAG (underlined) to eliminate RNA dimerization. The A* indicates the m^6^A modification site in the RNA fragment. **(C)** Binding of Y1-3 proteins to m^6^A RNA versus control, unmodified RNA was monitored using the AlphaScreen assay. Data are from two independent experiments with biological triplicates. Results are shown as mean ± SEM. ** P < 0.005 by Mann Whitney test. **(D)** Y1-3 protein pull-down using biotin-labeled control and m^6^A RNA fragments. Streptavidin Dynabeads were used in the pull-down assay and immunoblotting was performed using specific antibodies to Y1, Y2, or Y3. **(E)** Y1-3 protein pull-down level based on densitometry analysis of the bands in immunoblotting. Data are average results from two independent experiments. (**F**) Calculated concentrations of Y1-3 proteins needed to reach 50% pull-down levels of 625 nM Y1-3 by the m^6^A RNA fragment.

To study the binding properties of Y1-3 proteins to HIV-1 RNA with m^6^A modification, we synthesized two RNA fragments corresponding to nucleotides 235-281 of HIV-1_NL4-3_ gRNA with or without m^6^A modification in the second GGACU motif located in the 5’ UTR (Fig. 4A). To eliminate RNA dimerization in our binding assays, the underlined DIS sequence (AA<u>GCGCGC</u>) was replaced with the nucleotides GAG (Fig. 4A and 4B). We first used the AlphaScreen assay [20] to detect interaction of synthesized RNA fragments with purified full-length recombinant Y1-3 proteins. Each of these Y1-3 proteins exhibited clear preference for binding to m^6^A-modified HIV-1 RNA than its unmodified counterpart (Fig. 4C). We further investigated *in vitro* binding of the RNA fragments to Y1-3 protein using affinity pull-down experiments. Streptavidin-conjugated beads were used to pull-down biotin-modified control or m^6^A RNA fragments. Consistently, at lower Y1-3 concentrations (1-25 nM), clear preferences were seen for binding to m^6^A-modified HIV-1 RNA relative to control RNA, though both RNA fragments had detectable binding to Y1-3 proteins at higher concentrations (125-625 nM) (Fig. 4D). The immunoblotting results of the pull-down experiments were quantified and normalized to 1 based on protein pull-down levels by m^6^A RNA fragment at 625 nM protein input (Fig. 4E). To compare affinity of RNA fragments to Y1-3 proteins, we calculated the concentrations of each Y1-3 protein (ranging from 1-625 nM) required for 50% pull-down levels based on the regression curves (Fig. 4E and 4F). The 50% pull-down efficiencies indicated that Y1-3 proteins bound to m^6^A RNA fragment 7-fold, 13-fold, and >20-fold higher than control RNA, respectively (Fig. 4F). These results demonstrate that Y1-3 proteins exhibit substantially higher affinity for m^6^A-modified HIV-1 RNA *in vitro*, which may contribute to Y1-3-mediated inhibition of HIV-1 infection in cells.

### A to G mutations in GGACU motifs of the 5’ UTR of HIV-1 gRNA reduce viral infectivity

The two GGACU motifs are located in the PBS and an upstream region of the DIS (Fig. 4A), within the m^6^A peaks of the 5’ UTR of HIV-1 RNA that were identified in our previous study [13]. Because the 5’ UTR is critical for HIV-1 reverse transcription, genome package, and viral infectivity [21, 22], we further investigated the importance of these m^6^A sites on HIV-1 replication and infection by mutagenesis. To eliminate m^6^A modification of the GGACU motifs in HIV-1 gRNA, A to G mutations were introduced in the first (Mut1), second (Mut2), or both GGACT motifs (Mut3) in the HIV-1 proviral DNA plasmid pNL4-3 (Fig. 5A). WT pNL4-3 and derived mutants (Mut1-3) were separately transfected into HEK293T cells to measure HIV-1 protein expression and viral release. Compared with WT HIV-1, these mutants expressed comparable levels of precursor Gag protein, but the levels of cleaved p24 and intermediate Gag products in cell lysates were 1.4- to 1.8-fold higher (Fig. 5B). Consistently, supernatant p24 levels of mutant viruses were 1.2- to 1.4-fold higher than that of WT HIV-1 (Fig. 5C), suggesting a potential effect of these mutations on Gag proteolytic processing or viral release. To compare the infectivity of the mutants with WT HIV-1, viruses generated in HEK293T cells with equal amounts of p24 were used to infect TZM-bl indicator cells [23]. Interestingly, the infectivity of the mutant viruses was significantly lower (53-74%) relative to WT HIV-1 (Fig. 5D), suggesting an important role of these two GGACU motifs in HIV-1 infectivity.

**Fig. 5.**
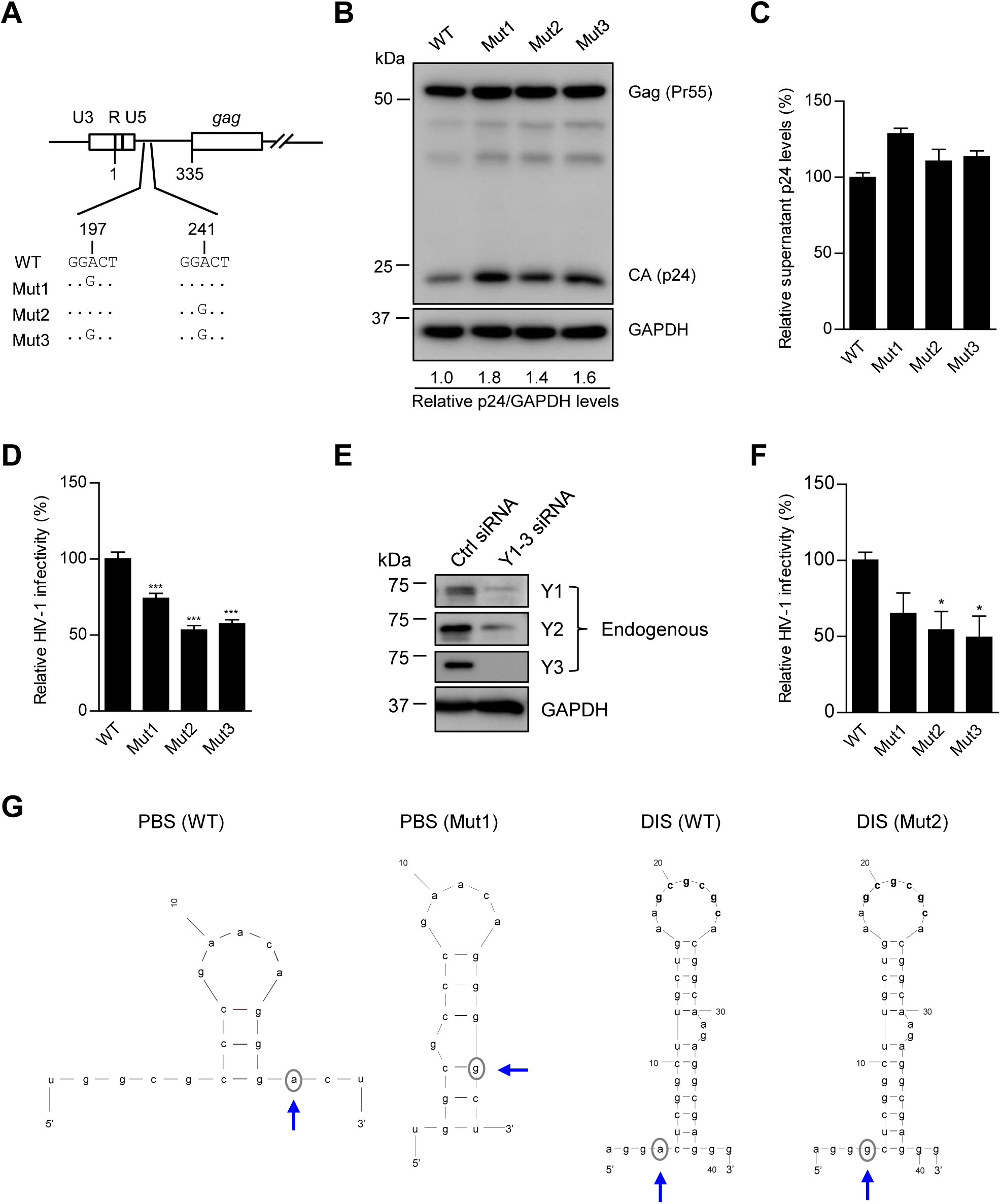
A to G mutations in GGACU motifs of the 5’ UTR of HIV-1 gRNA reduce viral infectivity. **(A)** Schematic representation of introduced mutations in the 5’ UTR region of HIV-1 proviral DNA. **(B)** Gag protein expression in HEK293T cell lysates expressing wild type (WT) and mutants (Mut1-3). The relative p24 levels were normalized based on GAPDH levels. **(C)** HIV-1 p24 levels in the supernatants at 48 h post-transfection were quantified by ELISA (n=3). **(D)** Infectivity of WT and mutant HIV-1 was quantified by measuring luciferase activity 48 h after infection of TZM-bl cells (n=4). Results are shown as mean ±SEM. Dunnett’s multiple comparison test was used for statistical analysis. *** P < 0.0005, compared HIV-1 mutants to WT virus. **(E)** Endogenous Y1-3 proteins in TZM-bl cells were knocked down by combined Y1-3 specific siRNA transfection. Immunoblotting confirms Y1-3 knockdown compared with control siRNA-transfected cells. **(F)** The infectivity of WT and mutant HIV-1 was quantified by measuring luciferase activity 48 h after infection of TZM-bl target cells (n=4). Results are shown as mean ±SEM. Dunnett’s multiple comparison test was used for statistical analysis. * P<0.05, compared HIV-1 mutants with WT HIV-1. **(G)** Predicted secondary structures of the RNA segments containing the PBS (left two panels) or DIS (right two panels, sequences are highlighted in bold) of WT, Mut1 and Mut2 viruses. Two A to G mutation sites (Mut1 and Mut2) are circled and pointed by blue arrows.

Previously published m^6^A mapping results indicate that there are multiple m^6^A sites in different regions of HIV-1 gRNA in addition to the 5’ UTR [11-13], which can potentially contribute to regulating viral infectivity through interactions with Y1-3 proteins. To address this, we examined whether silencing Y1-3 in target cells could restore the infectivity of these 5’ UTR mutant viruses. Endogenous Y1-3 proteins in TZM-bl cells were knocked down by combined Y1-3 specific siRNA, and then TZM-bl cells were infected with the same p24 amount of WT or mutant viruses generated from normal HEK293T cells. In TZM-bl target cells with partial Y1-3 proteins knockdown (Fig. 5E), all three mutant viruses showed 40-50% lower infectivity relative to WT HIV-1 (Fig. 5F). These results confirm the importance of these two GGACU motifs for HIV-1 infectivity, and suggest that other m^6^A sites in HIV-1 RNA can also regulate viral infectivity.

Given the important role of the PBS and DIS in structure and function of HIV-1 gRNA [24], we predicted the secondary structures of the RNA segments containing the PBS and DIS of WT and mutant viruses. Compared with the structure of WT HIV-1, the A to G mutation in the first GGACU motif (Mut1) resulted in a longer stem structure in the PBS sequence region, while mutation in the second GGACU motif did not change RNA structure containing the DIS (Fig. 5G). These results suggest that the decreased viral infectivity of Mut1 might be due to the combined effects of RNA structure change and elimination of m^6^A at this site. In contrast, the reduced viral infectivity of Mut2 was likely due to elimination of m^6^A modification.

### Y1-3 protein knockdown in virus-producing cells decreases HIV-1 Gag expression and alters viral production and infectivity

Our published results [13] and new data showed that Y1-3 proteins in target cells negatively regulate single-cycle and replication-competent HIV-1 infection (Fig. 1B and 2B-C, respectively). To further elucidate the role of Y1-3 in HIV-1 protein expression and infectivity, endogenous Y1-3 gene expression in virus-producing HEK293T cells was knocked down using specific siRNAs. Compared with non-specific control siRNA, single knockdown of each Y1-3 significantly decreased HIV-1 Gag (Pr55) protein expression, and correspondingly reduced the levels of processed p24 protein in cells (Fig. 6A). Consistently, p24 levels in the supernatants of cells with single or combined Y1-3 knockdown also reduced approximately 2-fold compared with those of control cells, and there was no synergistic effect of combined knockdown (Fig. 6B).

**Fig. 6.**
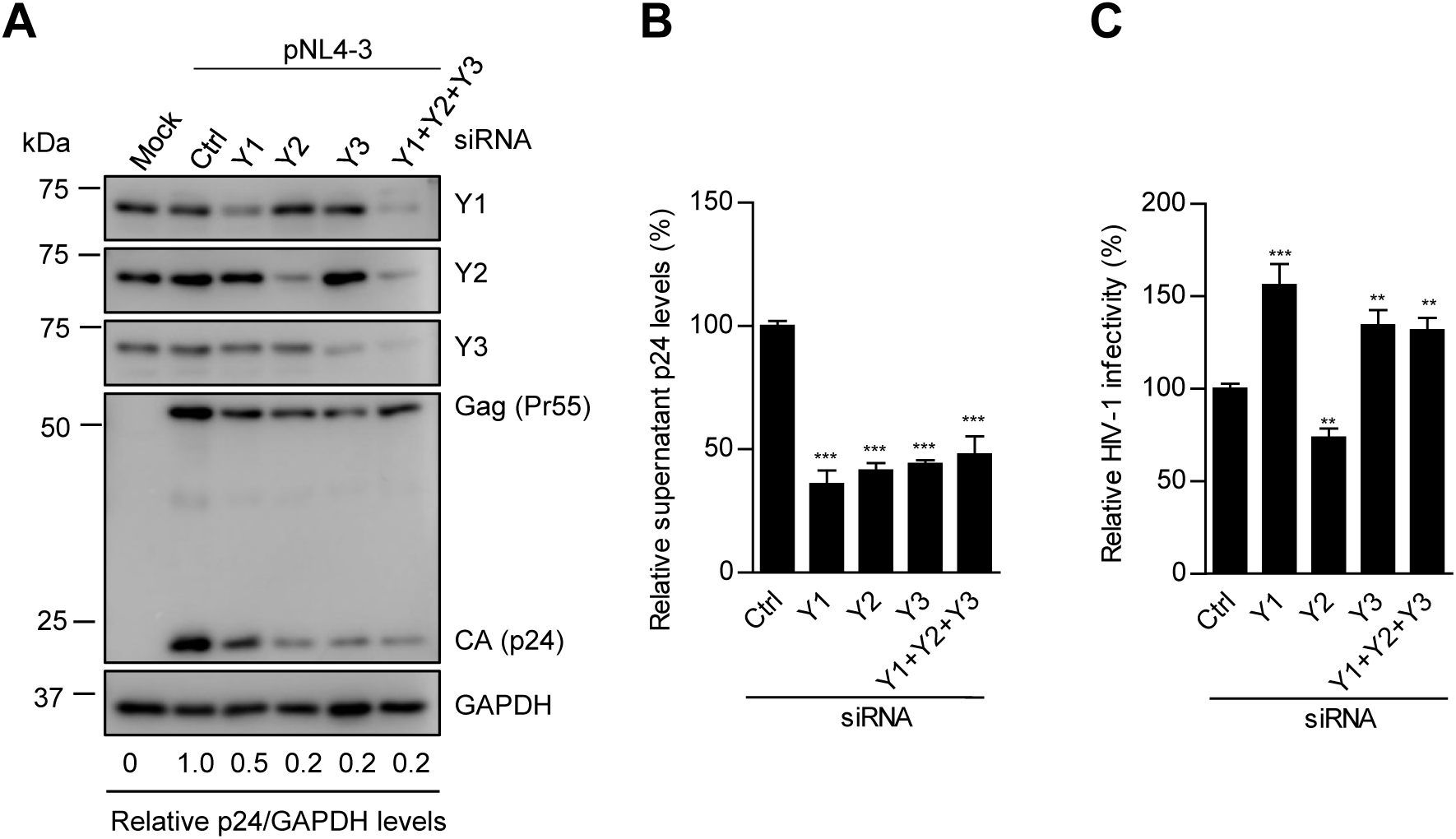
Knockdown of Y1-3 proteins in virus-producing cells affects HIV-1 Gag expression, viral production and infectivity. **(A)** Immunoblotting of HEK293T cells with siRNA-mediated single Y1-3 knockdown or Y1-3 combined knockdown and pNL4-3 transfection. (**B**) ELISA quantification of p24 levels in supernatants from the siRNA-transfected HEK293T cells (n=4). **(C)** Infectivity of HIV-1 generated in control cells, or cells with individual Y1-3 knockdown or combined knockdown was quantified by measuring luciferase activity 48 h after infection of TZM-bl reporter cells (n=3). Results are shown as mean ± SEM. Dunnett’s multiple comparison tests were used for statistical analysis. **P<0.005, *** P<0.0005, compared Y1-3 specific siRNA knockdown with non-specific control (Ctrl) siRNA.

To compare the infectivity of HIV-1 generated from cells with Y1-3 knockdown, viruses with equal amounts of p24 were used to infect TZM-bl indicator cells. As shown in Fig. 6C, HIV-1 generated from individual Y1 or Y3 knockdown cells had significantly higher infectivity compared with virus from control cells (P < 0.005), suggesting that Y1 and Y3 proteins in cells negatively affect infectivity of progeny HIV-1. In contrast, efficient Y2 knockdown in virus-producing cells resulted in a 25% decrease in HIV-1 infectivity (Fig. 6A and 6C), suggesting a different mechanism of Y2-mediated inhibitory effect on viral infection. Moreover, combined Y1-3 knockdown efficiently reduced endogenous levels of Y1-3 proteins in virus-producing cells (Fig. 6A, last lane), but only modestly increased HIV-1 infectivity (Fig. 6C), which were likely due to the different effects resulting from Y1/3 and Y2 knockdown.

### Y1-3 proteins and HIV-1 Gag form a complex with RNAs in HIV-1-infected target cells

To examine whether Y1-3 could interact with any HIV-1 proteins in cells, we performed IP of overexpressed Y1-3 in HEK293T cells co-transfected with pNL4-3, and then detected HIV-1 proteins in the input and IP products by immunoblotting using human anti-HIV-1 immunoglobulin [25]. Interestingly, we found that Y1-3 co-precipitated with HIV-1 Gag (Pr55) and intermediate Gag products, but not with HIV-1 p24 (Fig. 7, lanes labeled with IP). The vector cells were used as a negative control for IP and showed a background band of Gag (Pr55) in the IP product (Fig. 7, lane 2 from the left). Immunoblotting of FLAG confirmed expression and IP of FLAG-tagged Y1-3 proteins in the transfected cells. Because both Y1-3 proteins and HIV-1 Gag can bind cellular and HIV-1 RNAs [8, 10, 13, 26], to examine whether RNAs mediate the association between Gag and Y1-3 proteins, cell lysates were treated with RNase A before IP. Interestingly, RNase A treatment completely eliminated HIV-1 Gag signal in Y1-3 precipitation (Fig. 7, lanes labeled with RNase + IP). These results suggest that Y1-3 proteins and HIV-1 Gag form a complex with RNAs in cells.

**Fig. 7.**
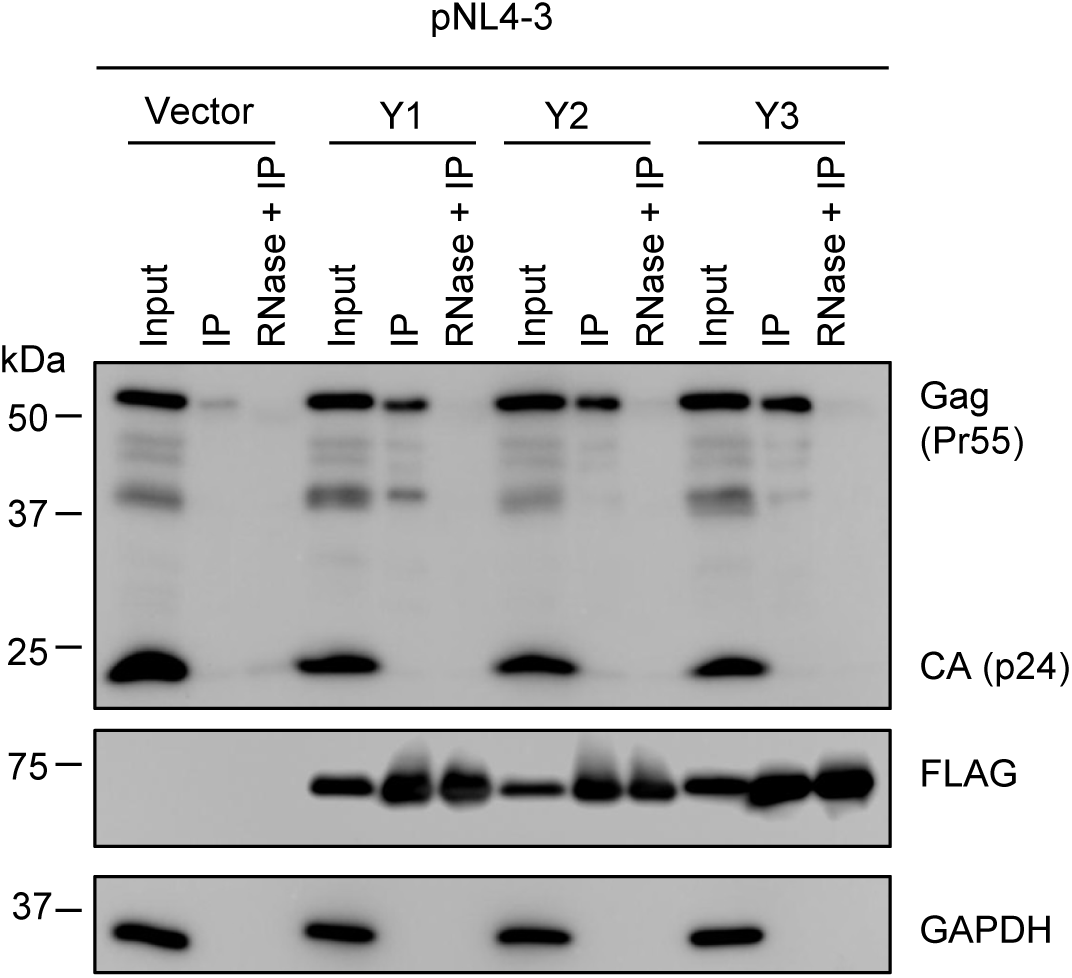
Y1-3 proteins and HIV-1 Gag form a complex with RNAs in HIV-1-infected target cells. HEK293T cells were co-transfected with pNL4-3 and individual plasmids expressing HA- and FLAG-tagged Y1-3 or empty vector. Cell lysates were treated with or without RNase A (100 µg/mL) before immunoprecipatation (IP) of Y1-3 proteins using anti-HA agarose beads. The input or IP products of HIV-1 Gag or CA (p24) were detected using HIV Immunoglobulin. Expression and IP of Y1-3 were confirmed with anti-FLAG immunoblotting. Results presented are representative of three independent experiments.

## Discussion

Reversible m^6^A modification is the most prevalent mRNA modification in eukaryotic organisms, and plays critical roles in gene expression [5]. Although m^6^A modification was previously identified in different viruses [27-29], its roles in HIV-1 gene expression regulation were recently recognized [11-13]. While these studies identified specific m^6^A sites in HIV-1 gRNA, they disagreed on the extent and locations of m^6^A modifications along the HIV-1 genome and their effects on viral replication [11-13]. Lichinchi *et al.* [11] reported 14 peaks of m^6^A modification in HIV-1 RNA, in which the m^6^A modification in the Rev response element region increased binding to Rev and facilitated nuclear export of viral RNA, thereby enhancing HIV-1 replication. In contrast, Kennedy *et al.* [12] found 4 clusters of m^6^A modification in the 3’ UTR region of HIV-1 RNA. Our study showed that HIV-1 gRNA has m^6^A modifications in the 5’ and 3’ UTRs as well as *gag* and *rev* genes [13]. Also, there is controversy regarding the roles of Y1-3 proteins in HIV-1 replication. In the presence of Y1-3 overexpression, Kennedy *et al*. observed increased HIV-1 replication [12], while our data showed decreased HIV-1 replication [13]. These discrepancies could stem from different cell types or lines, reagents, and techniques used, and need to be further investigated. In this study, we demonstrated that Y1-3 proteins suppress HIV-1 single-cycle and spreading infection in cells by decreasing viral gRNA and RT products. Y1-3 overexpression in HIV-1 infected cells led to significantly lower levels of early and late RT product, and further decreased HIV-1 infectivity, suggesting that Y1-3 proteins could also inhibit other steps after reverse transcription during the HIV-1 life cycle.

The GGACU is the predominant sequence in RRACH (R=G or A; H=A, C or U) motif recognized by the m^6^A writers [30]. Our *in vitro* biochemical experiments reveal that Y1-3 proteins exhibit clear preference for the m^6^A-modified HIV-1 RNA fragment over its unmodified RNA counterpart. The preferential binding of m^6^A sites in the HIV-1 genome to Y1-3 proteins may contribute to decreased HIV-1 infection in HeLa/Y1-3 cells. There are two GGACU motifs in the 5’ UTR of HIV-1_NL4-3_ gRNA. The first GGACU motif is in the PBS region, and is conserved in HIV-1 and simian immunodeficiency virus (SIV) [31], suggesting critical roles of the motif in HIV-1 and SIV life cycles. The second GGACU motif is in the DIS stem. This motif is conserved in HIV-1 subtype B isolates, but A to G mutations can be found in subtype C and group O isolates (https://www.hiv.lanl.gov/).

Compared with WT viruses, mutant viruses harboring A to G mutations have significantly reduced infectivity. We noted that the first GGACU motif is in the PBS and mutation at this site in Mut 1/3 viruses (A to G at nt. 197) introduces a mismatch with tRNA^lys^ primer [32], which may affect primer annealing and therefore impair viral infectivity. Recent studies showed that the second GGACU motif interacts with HIV-1 nucleocapsid (NC) protein in viral particles [32], and that nucleotides GGA in this motif strongly bind Gag precursor for viral genome packaging [33]. Mutation at this site may also impair interactions between HIV-1 gRNA and NC protein, contributing to decreased infectivity in mutant viruses. Thus, the decreased infectivity of Mut2 and Mut3 viruses (A to G at nt. 241) may be due to eliminated m^6^A modification at the second GGACU motif and/or decreased interactions between HIV-1 gRNA and Gag/NC. The effects of these mutations on primer binding and Gag/NC interactions remain to be examined.

Y1-3 proteins specifically recognizing m^6^A sites via a hydrophobic pocket in the YTH domain [3]. The effects of Y1-3 on HIV-1 gene expression, viral production in virus-producing HEK293T cells and infectivity of progeny HIV-1 appear complex. Individual or combined knockdown of each endogenous Y1-3 decreased HIV-1 Gag expression in cells and virion release in the supernatants (Fig. 6A and 6B). Knockdown of endogenous Y1 or Y3 proteins in virus-producing increased progeny HIV-1 infectivity, while Y2 knockdown modestly reduced the infectivity of progeny HIV-1 (Fig. 6C). These results together suggest that, in virus-producing cells, endogenous Y1-3 proteins positively regulate HIV-1 Gag protein expression, while Y1 and Y3 proteins negatively affect infectivity of the progeny HIV-1. Furthermore, the m^6^A modification regulates RNA processing and stability [2, 5, 34]. After HIV-1 transcription, viral RNA undergoes extensive splicing to express structural and accessory proteins. The roles of m^6^A modification in HIV-1 RNA splicing, stability and dimerization remain to be elucidated.

In addition to HIV-1, recent studies have identified viral RNA m^6^A modifications and its roles in regulating replication of *Flaviviridae* viruses, including hepatitis C virus (HCV), dengue, Zika virus (ZIKV), yellow fever virus, and West Nile virus [35, 36], Kaposi’s sarcoma-associated herpesvirus (KSHV) [37, 38], and influenza A virus (IAV) [39]. The m^6^A modification in viral RNA increases RNA expression of IAV [39], promotes KSHV lytic replication [37], but negatively regulates HCV and ZIKV production [35, 36]. Similar to our findings that Y1-3 proteins inhibit HIV-1 infection [13], Y1-3 proteins also negatively regulate HCV and ZIKV replication [35, 36]. A more recent study reported that Y2 protein negatively regulates the KSHV lytic replication by enhancing decay of KSHV transcripts [38]. Y1 and Y3 have no effect on IAV infection, while Y2 overexpression significant enhances IAV gene expression and replication [39]. Although different approaches in these studies may account for discrepancies, these results suggest that m^6^A modification and Y1-3 proteins have distinct effects on various viruses.

## Conclusions

We found that m^6^A reader proteins Y1-3 inhibit HIV-1 infection by decreasing viral gRNA and early reverse transcription products. We demonstrate that Y1-3 proteins preferentially bind to m^6^A-modified HIV-1 RNA. Mutation of m^6^A sites in the 5’ UTR resulted in decreased viral infectivity, suggesting important roles of these sites for HIV-1 infection. Y1-3 proteins and HIV-1 Gag form a complex with RNAs in virus-producing cells. Together, these results help better understand the roles and mechanisms of m^6^A modification of HIV-1 RNA in regulating viral replication.

## Methods

### Cell culture

HEK293T cell line (a kind gift from Dr. Vineet KewalRamani, National Cancer Institute, Frederick, USA) and TZM-bl cells [[23] obtained through the NIH AIDS Reagent Program, Catalog # 8129] were maintained in complete DMEM as described [40]. HeLa or HeLa/CD4 cell lines (kind gifts from Dr. Vineet KewalRamani, National Cancer Institute, Frederick, USA) overexpressing empty vector (pPB-CAG), individual FLAG- and HA-tagged Y1, Y2 or Y3 protein were maintained in complete DMEM containing 2 µg/mL puromycin as described [13].

### Generating HeLa cell lines stably express CD4 and Y1-3 proteins

HeLa/CD4 cells were generated by transduction of HeLa cells with a pMX retroviral vector expressing human CD4 [41]. HeLa/CD4 cells were transfected separately with pPB-CAG vector-based Y1-3 expressing constructs or pPB-CAG empty vector, and then selected with puromycin (2 µg/mL). HeLa/CD4 stably expressing cells Y1-3 were cultured as described [13].

### Mutagenesis in HIV-1 proviral DNA plasmid (pNL4-3)

Mutations in pNL4-3 were introduced using Agilent QuickChange Lightning Multi Site-directed Mutagenesis Kit (catalog # 210515-5) according to the instructions. Primers mut1 (5’-CCCGAACAGGG<u>G</u>CTTGAAAGC-3’) and mut2 (5’-TCGACGCAGG<u>G</u>CTCGGCTTG-3’) were used (mutation sites are underlined) to generate A to G mutation at the first (nt. 197), the second (nt. 241) or both GGACU motifs between HIV-1 gRNA U5 and *gag* gene. The mutations were confirmed by DNA sequencing.

### siRNA knockdown of Y1-3 proteins

Y1-3 expressions were knocked down by two rounds of siRNA transfection using specific siRNAs (QIAGEN) and Lipofectamine RNAiMax (Invitrogen) according to instructions (siRNA sequences are listed in Table 1). Briefly, HEK293T cells (1.5×10^5^ per well) in 24-well plates were transfected with gene specific siRNAs or control siRNA. Twenty-four h after transfection, cell culture media were replaced and the second-round siRNA transfection was conducted. At 6 h after the second round of siRNA transfection, pNL4-3 (0.5 µg per well) was transfected to HEK293T cells. Cells and culture media were collected at 36 h after pNL4-3 transfection for immunoblotting, p24 quantification, and infection assays.

**Table 1.**
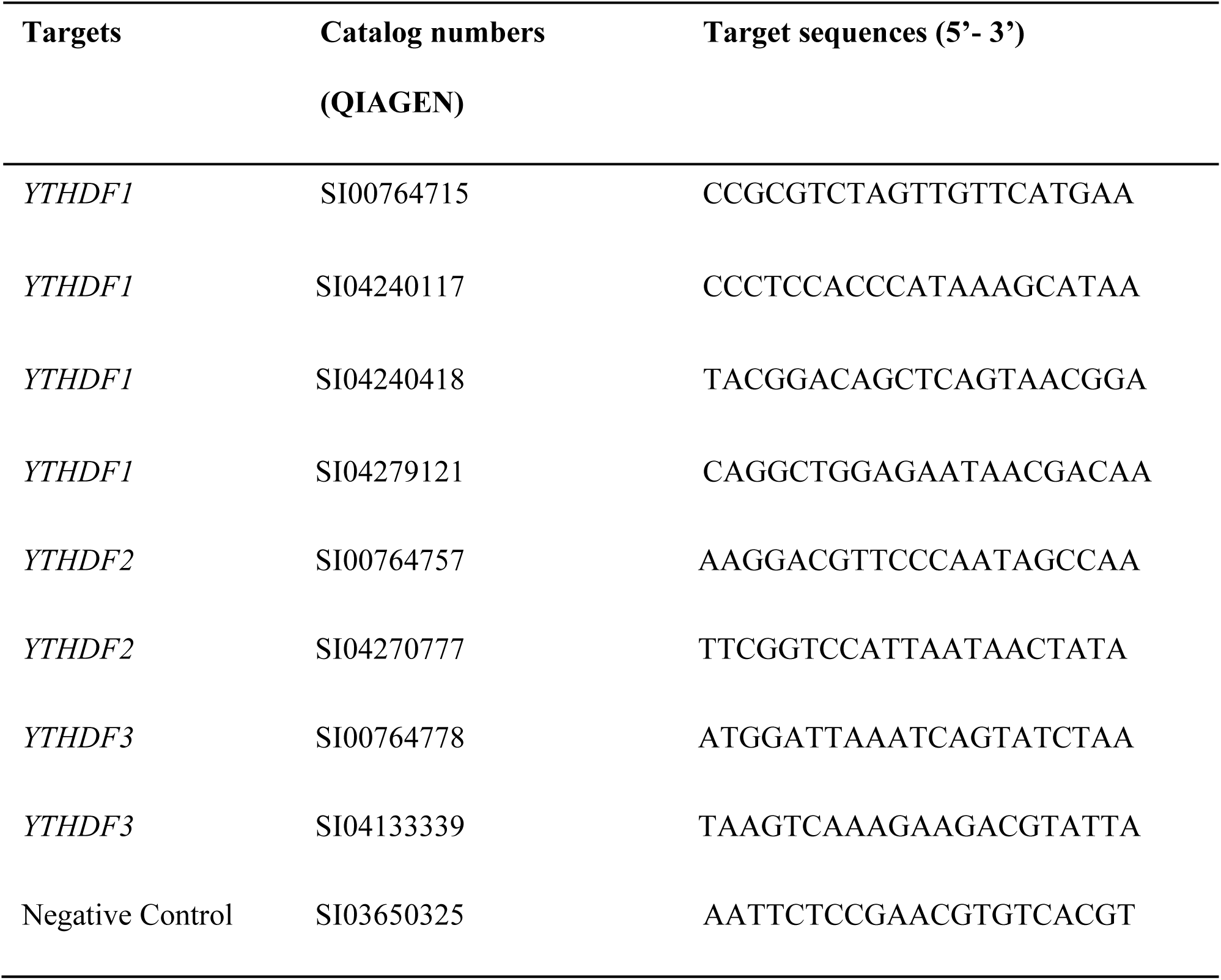
**Sequences of siRNAs used for Y1-3 knockdown**

### HIV-1 stocks and infection assays

Single-cycle HIV-luc/VSV-G was generated as previously described [13]. Replication-competent WT and mutant HIV-1 stocks were generated by transfection of HEK293T cells with pNL4-3 or mutant proviral DNA plasmids using Lipofectamine 2000 (Invitrogen) according to instructions. For Y1-3 and pNL4-3 co-expression experiments, pPB-CAG vector or pPB-CAG expressing individual FLAG- and HA-tagged Y1-3 protein was transfected into HEK293T cells, and pNL4-3 proviral DNA was transfected 12 h later. At 48 h (WT and Mut1-3 plasmids) or 36 h (Y1-3 co-expression) after proviral DNA transfection, cells were collected for immunoblotting and HIV-1 capsid p24 levels in viral stocks were quantified by an enzyme-linked immunosorbent assay (ELISA) using anti-p24-coated plates (AIDS and Cancer Virus Program, NCI-Frederick, MD) as described [13, 41]. To compare infectivity of WT HIV-1 in HeLa/Y1-3 cells, HIV-luc/VSV-G was used for infection at multiplicity of infection (MOI) at 1 as described [13]. To compare the infectivity of WT HIV-1_NL4-3_ and mutant viruses, viruses with equal amounts of p24 (400 pg) were used to infect TZM-bl cells in 24-well plates. At 48 hpi, TZM-bl cells were washed twice with PBS and lysed for luciferase assay (Promega) according to the manufacturer’s instructions. Cell protein concentrations were quantified using a bicinchoninic acid assay (Pierce) and all luciferase results were normalized based on total protein input.

### Flow cytometry measure cell surface levels of CD4 and CXCR4

HeLa/CD4 cells were treated with the non-enzymatic cell dissociation solution (C5914, Sigma-Aldrich) and double stained with FITC-conjugated CD4 (MHCCD0401-4, Thermo Fisher) and phycoerythrin-conjugated CXCR4 (555974, BD Pharmingen) antibodies as described [13]. Cells stained with a mouse IgG2a isotype antibody (555574, BD Pharmingen) were used as a negative control. Flow cytometry was analyzed using a Guava EasyCyte Mini and data analysis was performed using FlowJo (FlowJo, LLC) software as described [42].

### IP of Y1-3 proteins and RT-qPCR detection of HIV-1 gRNA

HeLa/CD4 cells expressing MAL were generated by transfection with pEF-Bos MAL Flag (plasmid # 41554 from Addgene) [17]. HeLa/CD4 cells expressing pPB-CAG vector, MAL or Y1-3 proteins (3 × 10^6^ cells) were seeded in a 60 mm-diameter culture plate one day before HIV-1 infection. Cells were infected with HIV-1_NL4-3_ at a multiplicity (MOI) of infection of 5 for 3 h. Cells were UV-cross-linked and lysed in cell lysis buffer (Sigma-Aldrich). Co-immunoprecipitated RNA was isolated and RT-qPCR quantification of HIV-1 gRNA was performed as described [13].

### Antibodies and immunoblotting

Antibodies used in this study were: anti-GAPDH (clone 4G5, AbD serotec, Atlanta, GA), anti-FLAG (F-3165, Sigma-Aldrich), anti-YTHDF1 (ab99080, Abcam), anti-YTHDF2 (ABE542, EMD Millipore, Billerica, MA), anti-YTHDF3 (ab103328, Abcam), and anti-HIV-1 p24 (AIDS Reagent Program, catalog # 6458). Cells were harvested and lysed in cell lysis buffer (Cell Signaling, Beverly, MA) supplemented with the protease inhibitor cocktail (Sigma-Aldrich). Immunoblotting was performed and ImageJ software (NIH) was used to calculate the densitometry of immunoblotting bands as described [13].

### AlphaScreen assay

Recombinant glutathione-S-transferase (GST)-tagged Y1-3 proteins were purified as described [8]. Two RNA fragments, control RNA and m^6^A-modified RNA (m^6^A RNA), corresponding to 235-281 nt containing the second GGACU consensus sequence, were synthesized (Integrated DNA Technologies), and the RNA fragments were modified with biotin at both 5’ and 3’ ends. Control RNA has no m^6^A modification, while m^6^A RNA fragment has m^6^A modification in GGACU motif. To eliminate RNA dimerization, the DIS sequence (AAGCGCGC) was substituted with nucleotides GAG as described in a previous study [43]. AlphaScreen assays were conducted as previously reported with minor modifications [20]. Proteins and RNA fragments were diluted with AlphaScreen buffer (100 mM NaCl, 1 mM MgCl_2_, 1 mM DTT, 1 mg/mL BSA, 25 mM Tris pH 7.5). For the binding of proteins and acceptor beads, Y1-3 proteins (25 nM) and acceptor beads (catalog # AL110C, 1:100 dilution; PerkinElmer) were mixed and adjusted to 30 µL, and incubated at 4 °C for 2 h. For RNA and donor bead binding, control or m^6^A RNA (50 nM) were mixed with donor beads (catalog # 6760002S, 1:100 dilution; PerkinElmer) in 10 µL volume, and incubated at 4 °C for 2 h. After the incubation, the protein and RNA were mixed, and incubated at 4 °C for 1 h. Samples (25 µL) were added to microplate and read with EnSpire multimode plate reader (PerkinElmer).

### *In vitro* pull-down assays for RNA and Y1-3 protein binding

Streptavidin Dynabeads M-280 (Invitrogen, catalog # 11205D) was used in this experiment. Beads were washed and incubated with biotin-labeled m^6^A RNA or control RNA at 4 °C for 60 min according to the manufacturer’s instructions, and purified Y1-3 proteins were added at concentrations of 1, 5, 25, 125, and 625 nM and incubated for 60 min. After washing, proteins bound to beads were eluted for immunoblotting using specific antibodies to Y1, Y2, or Y3.

### qPCR assays

To quantify HIV-1 gRNA after infection of HeLa/Y1-3 cells, at 1, 3 and 6 h after viral infection, cells were collected and total RNAs were extracted with an RNeasy Mini kit (Qiagen). Reverse transcription was conducted with first-strand synthesis (Invitrogen). qPCR was used to measure HIV-1 gRNA as described [44]. To quantify HIV-1 early and late RT products after viral infection, cellular DNA was extracted using a QIAamp DNA Blood Mini kit (Qiagen). Early RT products were quantified using primers ert2f (5’-GTGCCC GTCTGTTGTGTGAC-3’) and ert2r (5’-GGCGCCACTGCTAGAGATTT-3’). Late RT products were quantified using primers LW59 and LW60 as described [13]. GAPDH levels were also quantified to normalize early and late RT data [13].

### IP of Y1-3 proteins to detect the interactions with HIV-1 proteins

HEK293T cells (1 × 10^6^) were co-transfected with 2 µg pNL4-3 and 2 µg empty vector (pPB-CAG) or constructs expressing individual HA-tagged Y1-3 proteins as previously described [13]. At 48 h post-transfection, cells were harvested, lysed in 1% digitonin and total protein concentration was quantified. To elucidate the effects of RNAs in Gag and Y1-3 interactions, one aliquot of each cell lysate was treated with RNase A (100 µg/mL) for 1 h at room temperature, and IP was conducted as described [45]. Proteins bound to anti-HA agarose beads were eluted by boiling in sample buffer for immunoblotting. HIV Immunoglobulin (HIV-IG, NIH AIDS Reagent Program, catalog # 3957) [25] was used to detect HIV-1 proteins precipitated by Y1-3 proteins.

### Prediction of the secondary structure of HIV-1 RNA segments

The secondary RNA structures of HIV-1 5’UTR segments from WT HIV-1 or mutants were predicted using online mfold program according to the user’s instruction (http://unafold.rna.albany.edu/?q=mfold).

### Statistical analyses

Mann Whitney test was used to analyze AlphaScreen signal for m^6^A and control RNA fragment binding to Y1-3 proteins (Fig. 4C). Dunnett’s multiple comparison test was used for statistical analysis of all other data as indicated in figure legends. P <0.05 is considered significant.

## Acknowledgments

We thank Dr. Vineet KewalRamani for sharing reagents, Dr. Corine St. Gelais for critical reading the manuscript, and the Wu lab members for valuable discussions. The following reagents were obtained through the NIH AIDS Reagent Program, Division of AIDS, NIAID, NIH: HIV-1 p24 Gag Monoclonal (#24-3) from Dr. Michael H. Malim; HIV-IG (catalog #3957) from NABI and NHLBI; TZM-bl from Dr. John C. Kappes, Dr. Xiaoyun Wu and Tranzyme Inc.

## Ethics approval and consent to participate

Not applicable

## Consent for publication

Not applicable

## Author contributions

Conceived and designed the experiments: WL, NT, MK, LW. Performed the experiments: WL, NT, PCK. Analyzed the data: WL, NT, PCK, MK, CH, LW. Provided key reagents (purified recombinant Y1-3 proteins): CL. Wrote the paper: WL, NT, MK, LW.

